# Benchmarking Self-Supervised Learning for Single-Cell Data

**DOI:** 10.1101/2024.11.04.620867

**Authors:** Philip Toma, Olga Ovcharenko, Imant Daunhawer, Julia Vogt, Florian Barkmann, Valentina Boeva

**Affiliations:** Department of Computer Science, ETH Zurich

## Abstract

Self-supervised learning (SSL) has emerged as a powerful approach for learning biologically meaningful representations of single-cell data. To establish best practices in this domain, we present a comprehensive benchmark evaluating eight SSL methods across three downstream tasks and eight datasets, with various data augmentation strategies. Our results demonstrate that SimCLR and VICReg consistently outperform other methods across different tasks. Furthermore, we identify random masking as the most effective augmentation technique. This benchmark provides valuable insights into the application of SSL to single-cell data analysis, bridging the gap between SSL and single-cell biology.

## 1 Introduction

Recent progress in single-cell RNA sequencing (scRNA-seq) and multi-omics sequencing technologies have revolutionized our understanding of cellular heterogeneity and function at unprecedented resolution [1, 2]. While scRNA-seq measures gene expression levels in individual cells, generating a high-dimensional matrix where each row represents a cell and each column represents a gene’s expression level, multi-omics approaches expand this further by simultaneously profiling additional molecular layers such as the epigenome (ATAC-seq) or proteome (CITE-seq). A single multi-omics experiment can generate data matrices with hundreds of thousands of cells and multiple feature types - including expression of tens of thousands of genes from RNA-seq, hundred thousands of accessible chromatin regions from ATAC-seq, or hundreds of surface proteins from CITE-seq - creating an even higher-dimensional dataset where each cell is characterized by multiple measurements that collectively provide a more comprehensive view of cellular state and function. However, a major challenge in analyzing single-cell (multi-omics) data is the presence of batch effects—systematic variations arising from technical factors such as sample preparation, sequencing technologies, or laboratory conditions [3]. These batch effects can obscure the underlying biological signal and lead to wrong conclusions if not adequately addressed [4].

Successes of self-supervised learning (SSL) methods in computer vision [5, 6, 7], video processing [8], and natural language processing [9] have inspired their application to single-cell data. Several models have been adapted for analyzing single-cell data [10, 11, 12], showing promising results in mitigating batch effects and improving downstream analyses. However, a comprehensive benchmark comparing different SSL model architectures for single-cell data is currently lacking.

To address this need, we present a systematic benchmark of eight self-supervised learning methods applied to several large single-cell datasets. Our evaluation focuses on three downstream tasks: batch correction, query-to-reference mapping, and missing modality prediction [12]. In addition to evaluating model architectures, we compare several data augmentation techniques used for single-cell data. Data augmentation plays a crucial role in self-supervised learning by creating diverse views of the input data, enabling models to learn robust and invariant representations [6, 13].

**The main contributions** of our benchmark are (1) a systematic evaluation of eight SSL methods across eight different single-cell datasets, assessing their performance on three common downstream tasks; (2) evaluation of various hyperparameters, including representation and projection dimensionality, augmentation strategies, and multimodal integration methods, offering insights into optimal hyperparameter choices; (3) results that reveal SimCLR and VICReg outperform other architectures, with random masking constituting the best augmentation for data integration, both alone and combined with other transformations.

## 2 Benchmark Design

Self-supervised learning aims to learn useful data representations without relying on labels or other manual annotations [14]. Instead, SSL produces useful representations through representation invariance by leveraging the similarity and dissimilarity of data samples. For example, in contrastive SSL, different augmentations of the same image can be used to create positive pairs (i.e., similar examples), while distinct images can represent negative pairs (i.e., dissimilar examples) [6]. Alternatively, positive and negative pairs can comprise different modalities, such as image and text data [15]. Non-contrastive approaches leverage only positive pairs [16].

### Considered Methods

We benchmark eight existing SSL methods: SimCLR [6], MoCo [5], SimSiam [17], NNCLR [18], BYOL [7], VICReg [19], BarlowTwins [20], and Concerto [12]. See Figure 2 for details. First, in all methods except Concerto, two views are created by augmenting a single sample. Second, both views are encoded by a network with shared weights, producing data representations. Concerto removes the necessity for transforming samples by placing a dropout layer behind the encoder backbone. Finally, while training, all representations produced by the encoder are passed into a projector to improve robustness [21]. In all but Concerto, the projector is discarded during inference, keeping only the encoder’s output. SimCLR [6], MoCo [5], Concerto [12], and NNCLR [18] rely on positive and negative samples, unlike the other methods, which exploit negative-free learning. The emergence of non-contrastive methods was facilitated by an improved understanding of instabilities during model training. BYOL [7], NNCLR [18], and SimSiam [17] use a predictor to achieve better performance and avoid representation collapse, without leveraging negative pairs. Additionally, using momentum encoders in MoCo [5] and BYOL [7] helps against dimensionality collapse from, for instance, the lack of negative pairs. Finally, Concerto relies on a teacher-student network design to stabilize model training.

**Figure 1.**
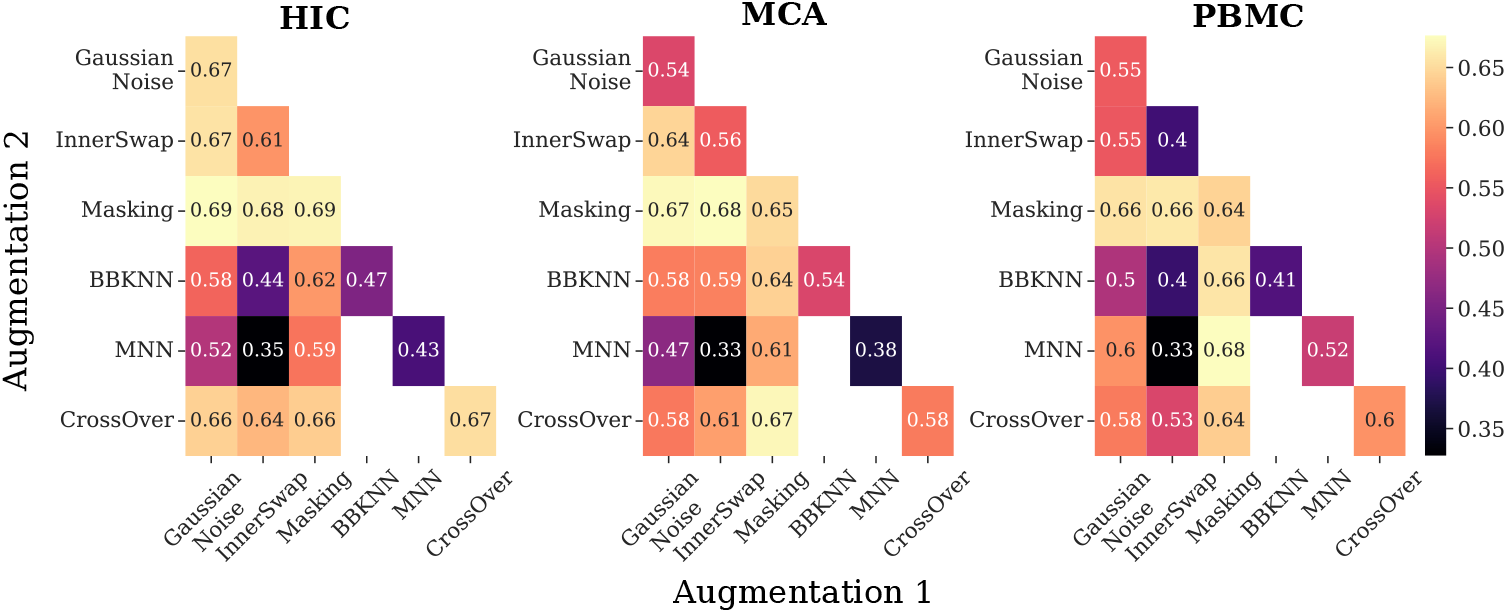
Evaluation of individual and combined data augmentations based on total score for batch correction. Diagonal entries correspond to a single augmentation, and off-diagonal entries correspond to the two sequentially applied augmentations. Hyperparameters based on ablation results (Table B5).

**Figure 2.**
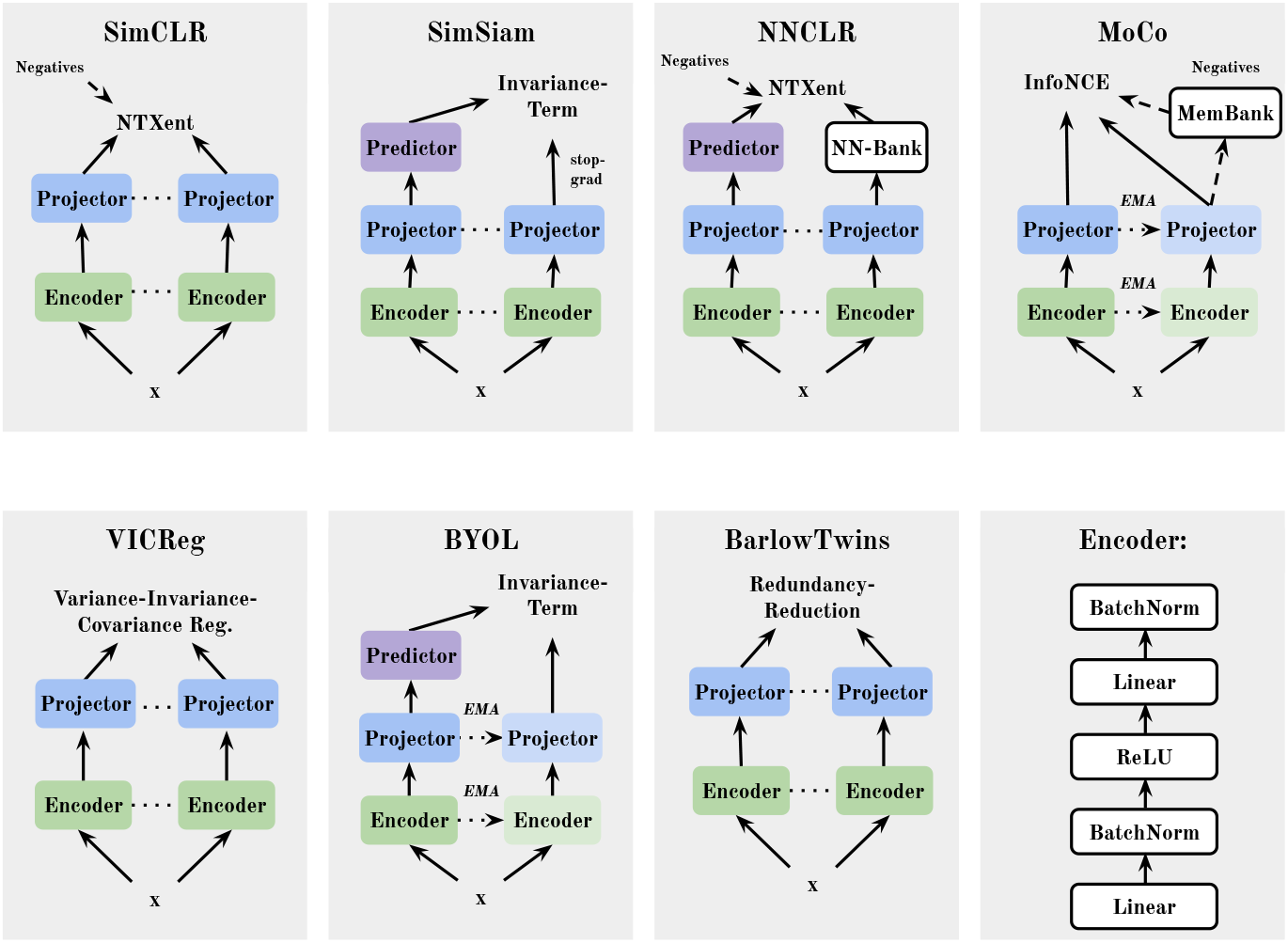
Overview of considered methods. Dotted lines between the encoder and projector blocks represent weight sharing. Exponential Moving Average (EMA) denotes the updating of weights with momentum. This figure was inspired by [5, 18, 19] based on our implementation of models with LightlySSL [35].

### Augmentations

We evaluate augmentations for single-cell data proposed in CLEAR [10] and CLAIRE [22]. The purpose of augmentations in SSL is to transform the original sample into two distinct views that are contrasted during training [23]. Multiple augmentations can be applied to a data sample for better generalization and robustness of representations. The authors of CLEAR [10] introduce four augmentations for scRNA-seq data that are each applied with 50% probability: Masking, Gaussian noise, InnerSwap, and CrossOver. First, a random mask sets 20% of a cell’s genes to zero, followed by additive Gaussian noise (with mean 0 and standard deviation 0.2) to 80% of genes in the cell. Then, 10% of genes are swapped within the cell (InnerSwap), before mutating 25% of gene expression values with another random cell (CrossOver). CLAIRE [22] uses a neighborhood-based approach when sampling cells for interpolation or mutation: mutual nearest neighbors in the unintegrated space are computed for each cell across all batches. During augmentation, an inter- and an intra-batch view are computed by mutating or interpolating between neighboring cells.

### Downstream Tasks and Evaluation

Our benchmark evaluates multiple single-cell datasets on three tasks: batch correction, query-to-reference mapping, and modality prediction.

Single-cell data can be affected by batch effects, which challenges the ability to measure true biological variation [24, 25]. Batch effects are technical biases introduced while sequencing because of differences in sequencing platforms, timing, reagents, or experimental conditions across laboratories [26]. To address batch effects, a common approach is learning a batch-corrected lower-dimensional embedding, where cells cluster based on their cell type rather than their experimental batch of origin [27]. The quality of batch-corrected embeddings is measured by biological conservation and batch correction metrics introduced in single-cell integration benchmarking (scIB) [28, 29], a tool that is widely used in the single-cell community, see subsection A.2 for more details. As introduced by Luecken et al. [29], we combine biological conservation (Bio) and batch correction (Batch) aggregate scores into a total score by *Total* = 0.6 *× Bio* + 0.4 *× Batch*.

Query-to-reference mapping is an unsupervised transfer learning task [12], where the primary objective is to annotate cells of a query dataset by mapping them to a joint latent space of a pre-annotated (reference) dataset [30]. Once query and reference data are aligned, cells of the query are annotated using a classifier trained on embeddings of the reference dataset (details in subsection A.2). k-nearest neighbor probing [31] is used to predict cell types, and performance is evaluated using the macro-average F1-score and classification accuracy [32].

For multimodal datasets, missing modality prediction enables the inference of unmeasured (missing) modalities in query cells [12]. Given a multimodal reference with RNA and protein expressions and a query containing only RNA, we predict the query’s original protein values by averaging reference proteins of the k nearest neighbors, see subsection A.2 for more details. We evaluate the quality of the inferred modality by measuring Pearson’s correlation between the original and predicted values.

## 3 Experiments

We benchmark eight self-supervised methods on eight single-cell datasets derived from different tissues with considerable variation in data size and complexity. Subsection A.1 provides more details about the datasets. All models are trained with five unique random seeds and reported with average performance and standard deviation.

### Hyperparameter tuning

We conducted hyperparameter tuning for all methods, except for Concerto, using two datasets: HIC and MAC. For Concerto, we use the hyperparameters proposed in [12]. For the other methods, we focus on two key hyperparameters: the representation and the projection dimensionality. First, we performed a grid search over the representation dimensionality for both datasets, evaluating the overall batch correction performance (details in subsection A.3). Our findings indicated that a dimensionality of 64 performed consistently well across all considered methods (see Figure 3). Expecting diminishing returns with further increases, we adopt this size for subsequent experiments. Additionally, we investigated the impact of projection dimensionality during training by introducing a scale factor. For contrastive methods, the projection size is scaled down by this factor, while for non-contrastive methods, it is scaled up by this factor (see subsection A.3). The results, presented in Figure 4, revealed that while the effect of the projector was ambiguous for most models, BarlowTwins, BYOL, and VICReg showed improved performance with larger up-scale factors.

**Figure 3.**
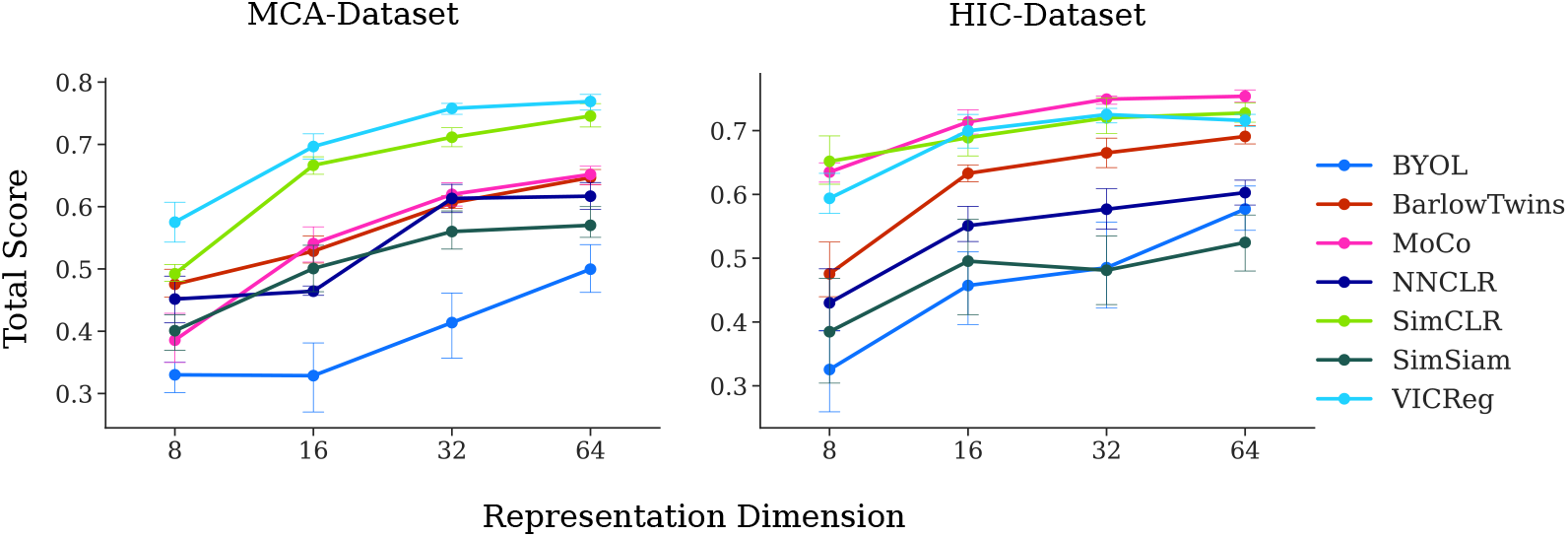
Tuning of the encoder based on the representation dimensionality. The encoder architecture is defined in subsection A.3. Lines correspond to the mean total score across five runs with unique seeds.

**Figure 4.**
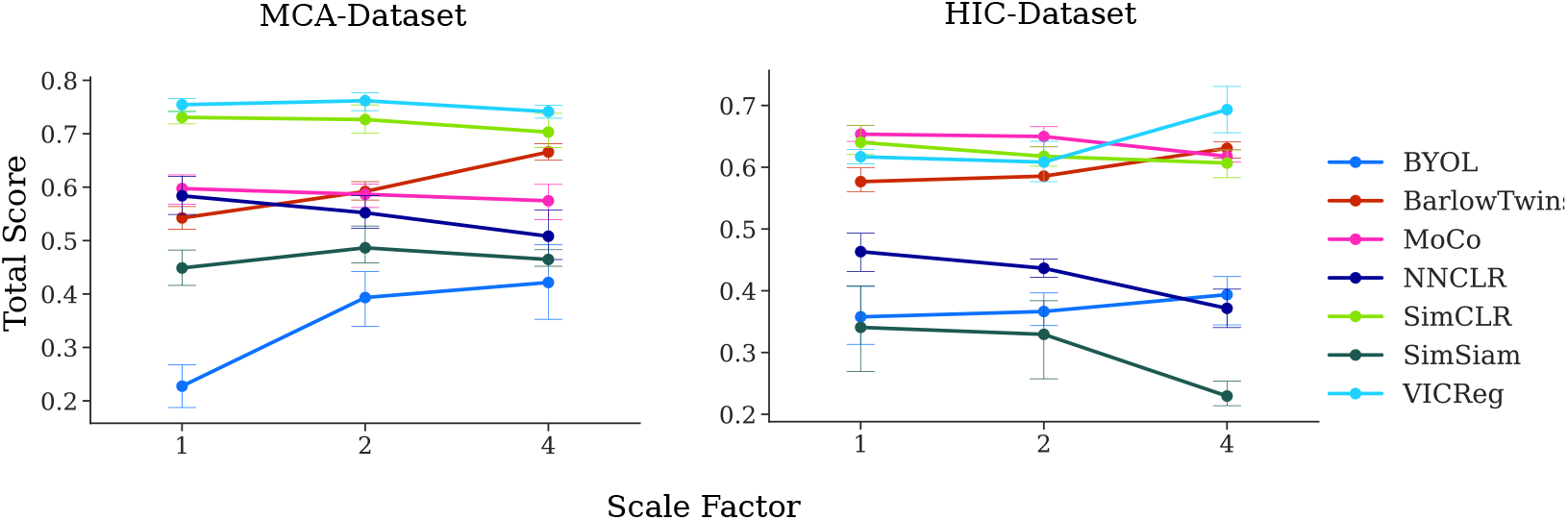
Tuning of the projector. The scale factor is defined in subsection A.3: in contrastive methods, the projected size decreased according to the scale factor, while for non-contrastive methods the projection size increases in accordance with the scale factor. Lines correspond to the mean total score across five runs with unique seeds.

### Batch correction

The batch correction performance of all methods across five datasets is presented in Table 1. Our analysis includes two multi-modal datasets, PBMC-CITE-seq and BMMC, along with three single-modality datasets. The results show that SimCLR outperforms all other architectures in terms of bio conservation across most datasets, while VICReg has a better batch correction and total score at the cost of bio conservation. Conversely, MoCo and BYOL overcorrect for batch effects, sacrificing biological signals, as can be observed on, e.g., the HIC and PBMC datasets.

**Table 1:**
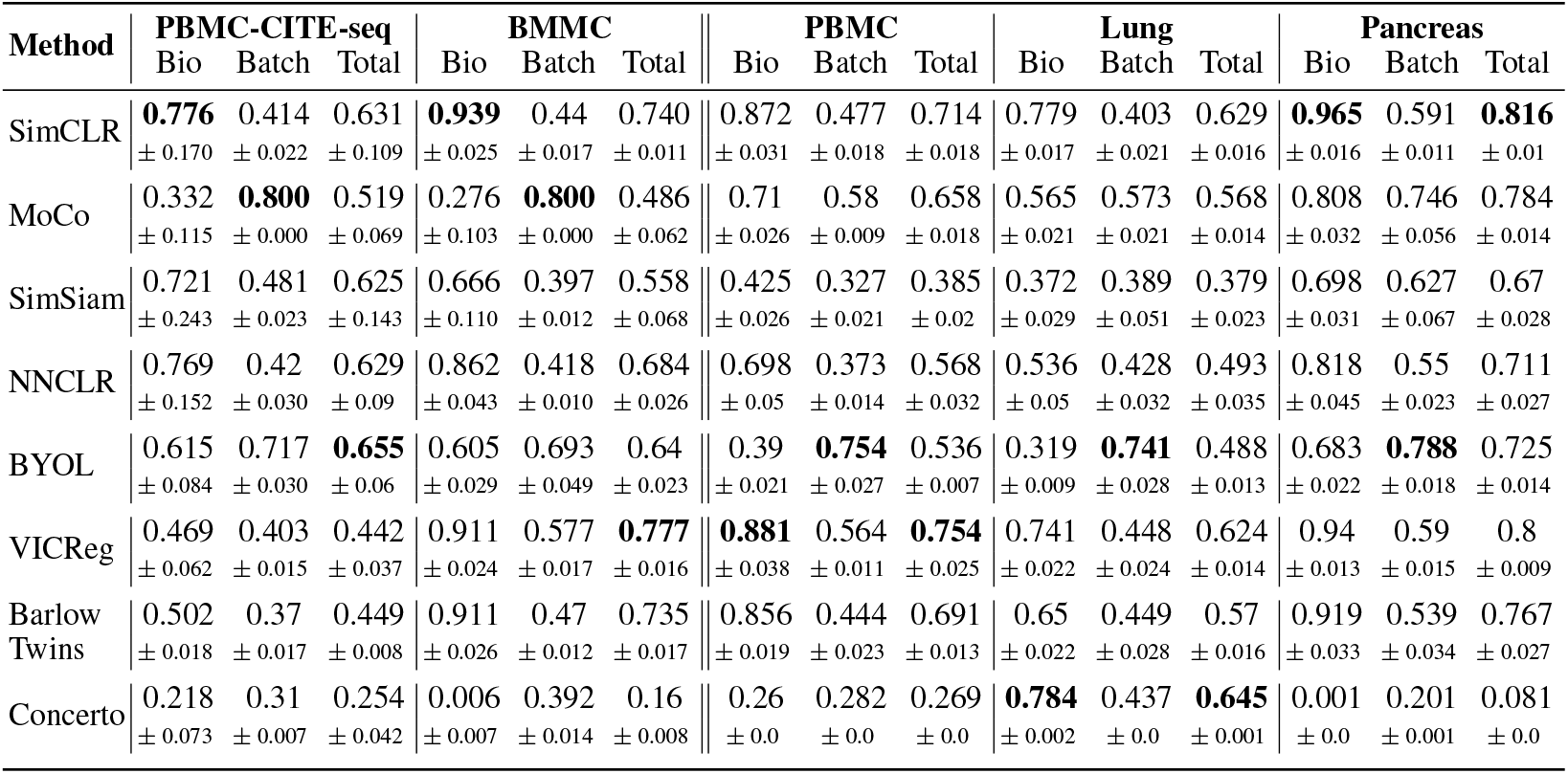
Batch integration performance across five datasets. Results show each method’s biological conservation score (Bio), batch correction score (Batch), and total score (Total), with means and standard deviations computed across five runs with different random seeds.

| Method | PBMC-CITE-seq |  |  | BMMC |  |  | PBMC |  |  | Lung |  |  | Pancreas |  |  |
| --- | --- | --- | --- | --- | --- | --- | --- | --- | --- | --- | --- | --- | --- | --- | --- |
|  | Bio | Batch | Total | Bio | Batch | Total | Bio | Batch | Total | Bio | Batch | Total | Bio | Batch | Total |
| SimCLR | **0.776**<br>± 0.170 | 0.414<br>± 0.022 | 0.631<br>± 0.109 | **0.939**<br>± 0.025 | 0.44<br>± 0.017 | 0.740<br>± 0.011 | 0.872<br>± 0.031 | 0.477<br>± 0.018 | 0.714<br>± 0.018 | 0.779<br>± 0.017 | 0.403<br>± 0.021 | 0.629<br>± 0.016 | **0.965**<br>± 0.016 | 0.591<br>± 0.011 | **0.816**<br>± 0.01 |
| MoCo | 0.332<br>± 0.115 | **0.800**<br>± 0.000 | 0.519<br>± 0.069 | 0.276<br>± 0.103 | **0.800**<br>± 0.000 | 0.486<br>± 0.062 | 0.71<br>± 0.026 | 0.58<br>± 0.009 | 0.658<br>± 0.018 | 0.565<br>± 0.021 | 0.573<br>± 0.021 | 0.568<br>± 0.014 | 0.808<br>± 0.032 | 0.746<br>± 0.056 | 0.784<br>± 0.014 |
| SimSiam | 0.721<br>± 0.243 | 0.481<br>± 0.023 | 0.625<br>± 0.143 | 0.666<br>± 0.110 | 0.397<br>± 0.012 | 0.558<br>± 0.068 | 0.425<br>± 0.026 | 0.327<br>± 0.021 | 0.385<br>± 0.02 | 0.372<br>± 0.029 | 0.389<br>± 0.051 | 0.379<br>± 0.023 | 0.698<br>± 0.031 | 0.627<br>± 0.067 | 0.67<br>± 0.028 |
| NNCLR | 0.769<br>± 0.152 | 0.42<br>± 0.030 | 0.629<br>± 0.09 | 0.862<br>± 0.043 | 0.418<br>± 0.010 | 0.684<br>± 0.026 | 0.698<br>± 0.05 | 0.373<br>± 0.014 | 0.568<br>± 0.032 | 0.536<br>± 0.05 | 0.428<br>± 0.032 | 0.493<br>± 0.035 | 0.818<br>± 0.045 | 0.55<br>± 0.023 | 0.711<br>± 0.027 |
| BYOL | 0.615<br>± 0.084 | 0.717<br>± 0.030 | **0.655**<br>± 0.06 | 0.605<br>± 0.029 | 0.693<br>± 0.049 | 0.64<br>± 0.023 | 0.39<br>± 0.021 | **0.754**<br>± 0.027 | 0.536<br>± 0.007 | 0.319<br>± 0.009 | **0.741**<br>± 0.028 | 0.488<br>± 0.013 | 0.683<br>± 0.022 | **0.788**<br>± 0.018 | 0.725<br>± 0.014 |
| VICReg | 0.469<br>± 0.062 | 0.403<br>± 0.015 | 0.442<br>± 0.037 | 0.911<br>± 0.024 | 0.577<br>± 0.017 | **0.777**<br>± 0.016 | **0.881**<br>± 0.038 | 0.564<br>± 0.011 | **0.754**<br>± 0.025 | 0.741<br>± 0.022 | 0.448<br>± 0.024 | 0.624<br>± 0.014 | 0.94<br>± 0.013 | 0.59<br>± 0.015 | 0.8<br>± 0.009 |
| Barlow<br>Twins | 0.502<br>± 0.018 | 0.37<br>± 0.017 | 0.449<br>± 0.008 | 0.911<br>± 0.026 | 0.47<br>± 0.012 | 0.735<br>± 0.017 | 0.856<br>± 0.019 | 0.444<br>± 0.023 | 0.691<br>± 0.013 | 0.65<br>± 0.022 | 0.449<br>± 0.028 | 0.57<br>± 0.016 | 0.919<br>± 0.033 | 0.539<br>± 0.034 | 0.767<br>± 0.027 |
| Concerto | 0.218<br>± 0.073 | 0.31<br>± 0.007 | 0.254<br>± 0.042 | 0.006<br>± 0.007 | 0.392<br>± 0.014 | 0.16<br>± 0.008 | 0.26<br>± 0.0 | 0.282<br>± 0.0 | 0.269<br>± 0.0 | **0.784**<br>± 0.002 | 0.437<br>± 0.0 | **0.645**<br>± 0.001 | 0.001<br>± 0.0 | 0.201<br>± 0.001 | 0.081<br>± 0.0 |

### Query-to-Reference Mapping

Table B1 and Table B2 show the query-to-reference performance for the single modal Pancreas ICA dataset with unique batches as queries. VICReg consistently outperforms all other models except for one query, with SimCLR showing similar accuracy and macro F1 scores. Table B3 evaluates multimodal query embeddings. We apply a concatenation to integrate two modalities and do not use a projector during inference. Additionally, we show the performance of a model trained on two modalities with CLEAR augmentations, but evaluated on a single main modality as a query, to verify whether representations contain information about the second modality. The second modality is unavailable during inference. Again, VICReg and SimCLR show the best performance, with VICReg performing slightly better. We observe that models trained and evaluated on multi-omics data perform better than the ones evaluated on a single modality. The performance drop is insignificant, and the model can be used for a single modality during inference if the second modality is missing.

### Missing Modality Inference

Table B4 shows the ability to predict missing protein values while given only RNA or gene expression (GEX) during inference. The model is trained on multi-omics data using CLEAR augmentations and concatenation to combine modalities. Again, SimCLR and VICReg perform better than the other methods. The high Pearson correlations show that models effectively infer protein values from gene expression data.

### Augmentation Ablation

The space of augmentations in the single-cell domain can be split into: random transformations [10, 33] and neighborhood-based transformations [11, 22]. We perform an ablation for all studied augmentations and optimize hyperparameters for each (see subsection A.4). To study how augmentations affect each other, we train a VICReg model using combinations of just two augmentations. We choose VICReg due to its consistently good performance. Figure 1 shows the best-performing augmentation is random masking, both alone and in combination with others.

### Comparison of Multimodal Integration Methods

In Table B6, we compare three methods to combine multiple modalities of a cell: element-wise addition of unimodal embeddings [12], concatenation of unimodal embeddings, and multimodal contrastive learning with the CLIP objective [15, 34]. For each modality, we train a modeland discard the projector during inference.See subsection A.5 for details. We use the CLEAR [10] pipeline for augmentation with the hyperparameters found in the single-modality ablation (Table B5). Table B6 shows that concatenation is the best-performing integration method overall. Both addition and concatenation show good results in bio conservation, while the CLIP-based approach performs better in batch correction.

## 4 Conclusions

We introduced a comprehensive benchmark for self-supervised learning with unimodal and multimodal single-cell data. We draw two main conclusions. First, we observe that the best-performing methods across all three downstream tasks are SimCLR and VICReg. Second, we conclude that masking augmentation leads to the biggest improvements alone and in combination with other types of augmentations. Interesting future work includes applying BBKNN augmentation for the multimodal experiments since it is to be explored how to apply it to multi-omics. Additionally, the inconsistent effects of projection during inference need further exploration.

## Code and Data Availability

The code to reproduce our results and preprocessed datasets are available at https://github.com/BoevaLab/scAugmentBench.

## Acknowledgments and Disclosure of Funding

We thank Sebastian Schelter and Sebastian Baunsgaard for their early feedback on the manuscript. Furthermore, we thank two anonymous reviewers for their thorough reviews, which significantly improved the quality of this paper.

FB is supported by the Swiss National Science Foundation (SNSF) (grant number 205321_207931).

## A Appendix

### A.1 Datasets

All datasets used in our benchmark are publicly available.

#### Human Immune Cells (HIC)

This dataset comprises 33,506 cells from 10 different donors assembled by Luecken et al. [29] from 5 studies. There are 16 cell types present in the data.

Availability: doi.org/10.6084/m9.figshare.12420968.v8

#### Mouse Cell Atlas (MCA)

This dataset comprises 6954 cells collected across two studies [36]. Three different sequencing protocols were used, and 11 cell types are present in the dataset.

Availity: figshare.com/10351110 and figshare.com/10760158

#### Peripheral Blood Mononuclear Cells (PBMC)

Collected by Ding et al [37], this dataset contain 30,449 cells from 2 patients. Cells were sequenced with seven different protocols.

Availability: singlecell.broadinstitute.org/single-cell-comparison-pbmc-data

#### Pancreas

This dataset was collected by Tran et al. [36] combining five studies of the human pancreas. It comprises 14,767 cells sequenced by one of four scRNA-seq technologies.

Availability: figshare.com/24539828

#### Lung

This dataset contains 32,426 cells across 16 batches and two technologies, assembled by Luecken et al. [29]. The cells are from human peripheral blood and human bone marrow.

Availability: figshare.com/24539942

#### Immune Cell Atlas

This dataset contains 329,762 cells across 12 batches and three different technologies, collected by Conde et al. [38]. The cells originate from 16 different tissues.

Availability: cellxgene.cziscience.com/08f58b32-a01b-4300-8ebc-2b93c18f26f7

#### Multimodal Peripheral Blood Mononuclear Cells (PBMC CITE-seq)

This dataset was collected by Hao et al. [39] with 161,764 cells across 8 batches. For each cell, two modalities are available: RNA and protein. As a pre-processing step, we merge different T cell granularities, similar to the Concerto framework [12], and remove cells annotated as *other* to reduce noise.

Availability: atlas.fredhutch.org/pbmc_multimodal.h5seurat

#### Multimodal Bone Marrow Mononuclear Cells (BMMC)

This dataset was collected by Luecken et al. [40] and contains 90,261 cells across 13 batches and 12 healthy human donors [41]. Each cell has two modalities: Gene expression (GEX) and protein abundance (ADT). Pre-processing is the same as PBMC CITE-seq.

Availability: ncbi.nlm.nih.gov/acc.cgi?acc=GSE194122

#### A.2 Evaluation Details

##### Preprocessing

All datasets are preprocessed using SCANPY [42] normalize-total function, which scales the total counts per cell to 10,000, followed by log-transformation. We subsequently perform batch-aware feature selection to choose the 4,000 most highly-variable genes (HVGs) for further processing. For multimodal PBMC CITE-seq and BMMC datasets, we select 2,000 HVGs contrary to 4,000 HVGs for the single modality datasets.

##### Batch Correction

The evaluated metrics are divided into two categories: those that measure the conservation of biological variance and that that measure the batch correction [29, 36]. To evaluate conservation of biological variation, we calculate the isolated labels score, the Leiden NMI and ARI, the silhouette label score, and the cLISI metric. To evaluate batch correction, we calculate the graph connectivity, kBET per label, iLISI for each cell, the PCR comparison score, and the silhouette coefficient per batch. For details and definitions of the used evaluation metrics, as well as their implementation, we refer to [29]. All tables showing batch correction results are min-max scaled inside each dataset, and each method’s evaluation metric is scaled individually before aggregating scores.

##### Query-to-Reference Mapping and Missing Modality Prediction

In the PBMC CITE-seq dataset, for query-to-reference mapping and missing modality inference, we hold out batches *P3, P5*, and *P8*. In the BMMC dataset, for query-to-reference mapping and missing modality inference, we hold out batches *s4d1, s4d8*, and *s4d9*. Similar to the approach of [43], we perform query-to-reference mapping by fitting a non-parametric supervised classifier (k-nearest neighbors (kNN) classifier with *k* = 11). For missing modality prediction, we fit a kNN classifier with *k* = 5, as in [12].

#### A.3 Hyperparameter Tuning

In all experiments, we use the augmentation pipeline proposed by CLEAR [10] as a foundation, unless stated differently. Experiments described in this section were computed for all methods except Concerto. For the latter, we use the original model from [12].

##### Optimization

All models in this benchmark, except Concerto, were trained with the Adam optimizer [44]. We use a stepwise learning rate schedule with base learning rate 1e-4 and fix the batch size at 256. When applicable, the memory bank size was set to 2048.

##### Encoder Architecture

We fix the encoder across all architectures and only perform a hyperparameter search on the dimensionality of the encoder output, i.e., the representation dimensionality. The encoder consists of a fully connected layer reducing the dimensionality to 128, followed by a ReLU activation and batch normalization. A further fully connected layer encodes the hidden representation to the representation dimension, followed by batch normalization.

##### Representation Dimensionality

We search for the best representation dimensionality by training all models with dimension {8, 16, 32, 64} across five runs with different random seeds. Models are ranked according to the SCIB-METRICS total score, which is min-max scaled across all model instances.

##### Projector Dimensionality

Projection heads benefit self-supervised models in learning robust representations [45]. At inference, the projection head is discarded, and only the (backbone) encoder is used for inference. All evaluated architectures subject to our evaluation include a projection head. We perform a hyperparameter search to find the best output dimension of the projector.

All projection heads were implemented as noted in the respective works. In their respective works, SimCLR, MoCo, SimSiam, and NNCLR are evaluated with projectors that retain or scale down the dimensionality of the representation. BarlowTwins, BYOL, and VICReg are evaluated with projectors that retain or scale up the dimensionality. We follow this rationale and search a grid of scaling factors {1, 2, 4}. To compute the projection dimensionality, the scaling factor is either divided (scale-down models) or multiplied (scale-up models) with the representation’s dimension.

##### Regularization Hyperparameters

Variance-invariance-covariance regularization hyperparameters are used as is done in the original work. We evaluate a grid of parameters, where the invariance term and the variance term *λ, α* = {5, 10, 25, 50}, while the invariance term *β* is fixed to 1. We find that *λ* and *α* fixed to 5 perform well across both ablation datasets.

##### Augmentation Strength

Augmentations are known to benefit SSL models in finding robust representations. Details of the evaluated augmentations are listed in subsection A.4. We perform a grid search to optimize the hyperparameters for all augmentations. This includes *α* for all models, *σ* for the Gaussian Noise augmentation, and the *kNN*-size for the nearest-neighbor-based transforms MNN and BBKNN. For each augmentation, the original CLEAR hyperparameters are fixed, and only the hyperparameters of the evaluated augmentation are adapted. For the ablation of BBKNN, we remove CrossOver, and replace it by BBKNN. Due to the implementation of MNN, we remove CrossOver and insert MNN at the front of the augmentation pipeline. Results of the ablation are recorded in Figure 5.

**Figure 5.**
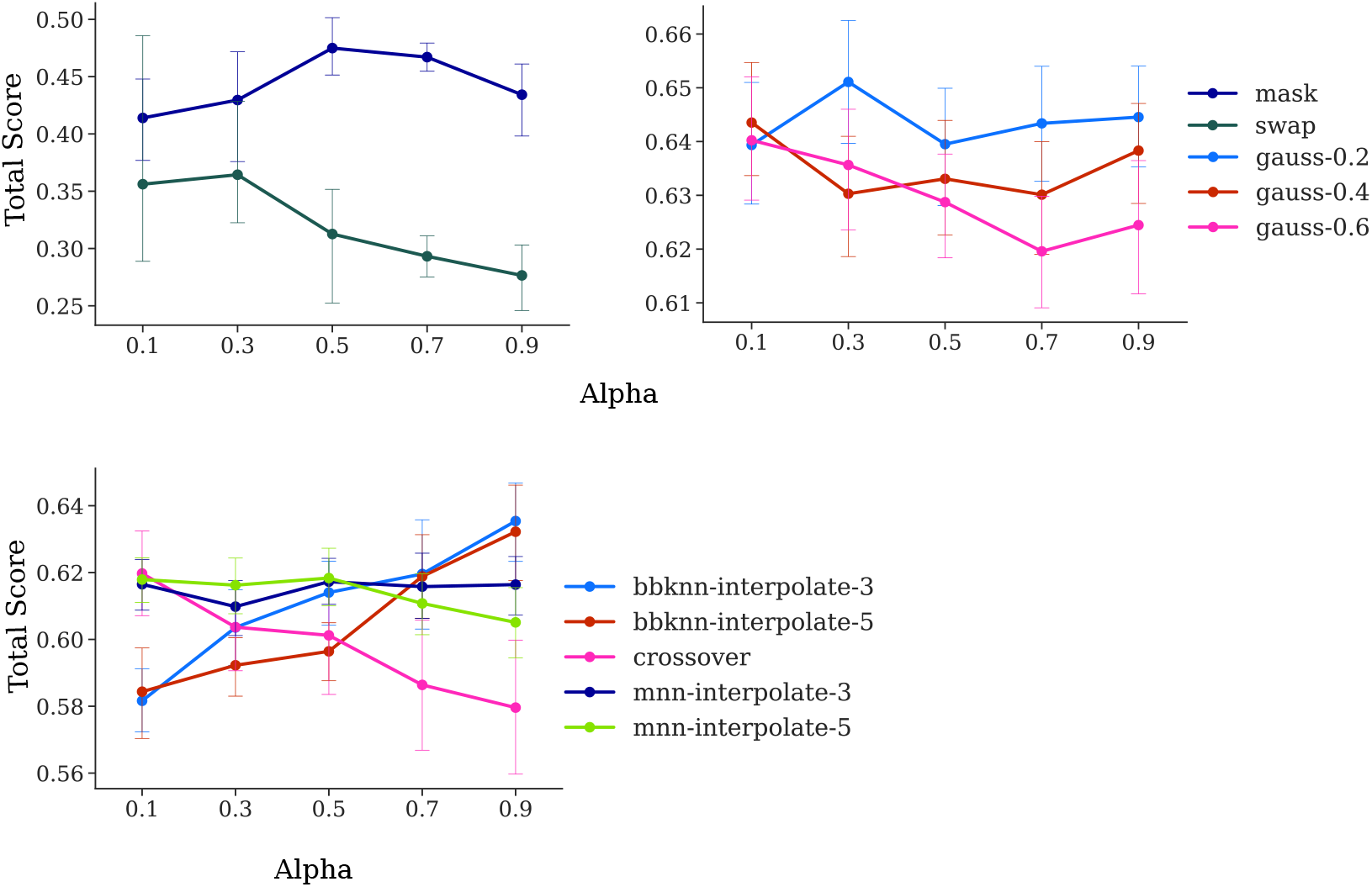
Ablation on the augmentation hyperparameters. The figure aggregates results for all methods, trained on the HIC dataset.

#### A.4 Augmentations

We evaluate six augmentations in this work. For all, the parameter *α* defines the proportion of values affected by the transform. Augmentations are applied sequentially. Masking is performed by setting gene expressions to zero. Gaussian noise computes a noise vector computed from the normal distribution (with zero mean and standard deviation *σ*) and adds it to the input. InnerSwap switches expressions between genes within a cell, while CrossOver switches expressions of the same gene between two *random* cells.

The MNN augmentation refers to our implementation of CLAIRE’s augmentation [22]. For each cell, it computes an intra- and inter-batch neighborhood based on its mutual nearest neighbors. Then, views are computed by interpolating between neighbors. We do not filter cell-neighborhoods based on representation similarities during early stages of model training, as is done in the original work. This work introduces the BBKNN augmentation. It uses a non-trimmed batch-balanced kNN graph [25] to define a set of *#batches* × *knn* neighbors for each cell. Views are computed by interpolating between neighbors. It differs from CLAIRE’s concept in that it does not distinguish between intra- and inter-batch neighbors. While MNN always produces a view based on neighbors within and a view based on neighbors from outside the batch, this is not the case for BBKNN. Due to its implementation, the MNN augmentation is limited to be applied first in any augmentation pipeline. We refer to [22] for further detail on the interpolation process.

#### A.5 Multimodal Setting

Recent developments in the single-cell analysis allow the measurement of multiple aspects of a cellular state. Data containing multiple modalities of a cell, e.g., RNA and protein, is called multiomics. Existing self-supervised methods for single-cell data integration can be extended to the multimodal setting by combining views produced by specialized models for different modalities. We train two models for each modality; each model consists of an encoder and projector. As is common [5, 6, 14], only the encoder is used to infer the integrated representation. However, in the single-cell community, the projector is also used during inference, and, therefore, we also evaluate whether projection during prediction improves performance. Additionally, there are various techniques to combine representations [12, 15, 34]. We evaluate three approaches: Addition, concatenation, and CLIP.

##### Three Multimodal Integration Methods

First, **addition** takes two embeddings of two modalities and adds them together to get a joint representation, similar to the Concerto framework [12]. A constraint is that both embeddings should be of the same dimensionality. Second, **concatenation** appends two embeddings (not necessarily of the same size). Third, instead of contrasting two joint views of a cell, two modalities of the same cell are contrasted using symmetric cross-entropy loss [46] and **CLIP approach**. After training leveraging the **CLIP approach** [15, 34], we concatenate two embeddings during inference.

##### Encoder & Projector Embedding Evaluation

Using CLEAR augmentations, we train two models for each modality, each consisting of an encoder and projector. In Table B7, we compare data integration performance with and without a projector during inference. Interestingly, SimCLR benefits from projection, while VICReg performance degrades. We conclude that the effect of projection is inconsistent across models.

### B Supplementary Tables

**Table B1:** Query-to-Reference Mapping with CLEAR augmentations on the Pancreas dataset. We define individual studies as holdout sets during training. Accuracy and Macro F1 are computed on the holdout set.

| Method | Mutaro et al. |  | Segerstolpe et al. |  | Wang et al. |  | Xin et al. |  |
| --- | --- | --- | --- | --- | --- | --- | --- | --- |
|  | Macro F1 | Acc | Macro F1 | Acc | Macro F1 | Acc | Macro F1 | Acc |
| SimCLR | **0.701** | **0.963** | 0.677 | 0.975 | 0.665 | 0.929 | 0.304 | 0.783 |
| | $\pm 0.052$ | $\pm 0.002$ | $\pm 0.027$ | $\pm 0.003$ | $\pm 0.046$ | $\pm 0.006$ | $\pm 0.031$ | $\pm 0.009$ |
| MoCo | 0.653 | 0.962 | 0.68 | 0.974 | 0.678 | 0.928 | 0.307 | 0.771 |
| | $\pm 0.04$ | $\pm 0.002$ | $\pm 0.02$ | $\pm 0.003$ | $\pm 0.01$ | $\pm 0.012$ | $\pm 0.057$ | $\pm 0.012$ |
| SimSiam | 0.589 | 0.922 | 0.615 | 0.935 | 0.592 | 0.887 | 0.225 | 0.701 |
| | $\pm 0.019$ | $\pm 0.016$ | $\pm 0.036$ | $\pm 0.014$ | $\pm 0.035$ | $\pm 0.016$ | $\pm 0.039$ | $\pm 0.013$ |
| NNCLR | 0.674 | 0.952 | 0.631 | 0.941 | 0.641 | 0.916 | 0.246 | 0.729 |
| | $\pm 0.054$ | $\pm 0.008$ | $\pm 0.028$ | $\pm 0.009$ | $\pm 0.05$ | $\pm 0.015$ | $\pm 0.034$ | $\pm 0.011$ |
| BYOL | 0.611 | 0.945 | 0.639 | 0.942 | 0.597 | 0.867 | 0.274 | 0.68 |
| | $\pm 0.021$ | $\pm 0.003$ | $\pm 0.009$ | $\pm 0.005$ | $\pm 0.035$ | $\pm 0.023$ | $\pm 0.045$ | $\pm 0.025$ |
| VICReg | 0.652 | 0.961 | **0.689** | **0.978** | **0.687** | **0.937** | **0.339** | **0.81** |
| | $\pm 0.036$ | $\pm 0.002$ | $\pm 0.037$ | $\pm 0.003$ | $\pm 0.035$ | $\pm 0.003$ | $\pm 0.042$ | $\pm 0.007$ |
| Barlow Twins | 0.629 | 0.947 | 0.62 | 0.928 | 0.624 | 0.905 | 0.2 | 0.69 |
| | $\pm 0.053$ | $\pm 0.005$ | $\pm 0.03$ | $\pm 0.01$ | $\pm 0.025$ | $\pm 0.011$ | $\pm 0.014$ | $\pm 0.012$ |
| Concerto | 0.106 | 0.431 | 0.113 | 0.419 | 0.105 | 0.435 | 0.112 | 0.406 |
| | $\pm 0.0$ | $\pm 0.0$ | $\pm 0.0$ | $\pm 0.0$ | $\pm 0.0$ | $\pm 0.0$ | $\pm 0.0$ | $\pm 0.0$ |

**Table B2:** Unimodal Query-to-Reference Mapping with CLEAR augmentations. We define one technology (10X 5’ v2) of the Immune Cell Atlas as a holdout set, train the encoder and knn-classifier, and evaluate performance on the holdout set.

|  | Method |  |  |  |  |  |  |
| --- | --- | --- | --- | --- | --- | --- | --- |
|  | SimCLR | MoCo | SimSiam | NNCLR | BYOL | VICReg | Barlow Twins |
| Macro F1 | 0.788 | 0.794 | 0.711 | 0.74 | 0.68 | **0.82** | 0.727 |
| | $\pm 0.004$ | $\pm 0.006$ | $\pm 0.015$ | $\pm 0.01$ | $\pm 0.002$ | $\pm 0.012$ | $\pm 0.004$ |
| Acc | 0.83 | 0.835 | 0.768 | 0.804 | 0.724 | **0.866** | 0.752 |
| | $\pm 0.001$ | $\pm 0.014$ | $\pm 0.007$ | $\pm 0.007$ | $\pm 0.009$ | $\pm 0.003$ | $\pm 0.007$ |

**Table B3:** Query-to-reference for multimodal datasets with CLEAR pipeline. On the left, two modalities (RNA + Protein or GEX (gene expression) + ADT (protein abundance)) were used during inference. On the right, we show inference performance with a single modality (RNA or GEX). All models were trained with two modalities.

| Method | RNA + Protein |  | GEX + ADT |  | RNA |  | GEX |  |
| --- | --- | --- | --- | --- | --- | --- | --- | --- |
|  | PBMC CITE-seq<br>Macro F1 | Acc | BMMC<br>Macro F1 | Acc | PBMC CITE-seq<br>Macro F1 | Acc | BMMC<br>Macro F1 | Acc |
| SimCLR | **0.989**<br>$\pm 0.001$ | **0.994**<br>$\pm 0.001$ | 0.839<br>$\pm 0.009$ | 0.894<br>$\pm 0.005$ | 0.958<br>$\pm 0.004$ | **0.957**<br>$\pm 0.004$ | 0.843<br>$\pm 0.003$ | 0.877<br>$\pm 0.006$ |
| MoCo | 0.94<br>$\pm 0.006$ | 0.951<br>$\pm 0.006$ | 0.725<br>$\pm 0.029$ | 0.802<br>$\pm 0.028$ | 0.84<br>$\pm 0.014$ | 0.838<br>$\pm 0.012$ | 0.7<br>$\pm 0.027$ | 0.717<br>$\pm 0.033$ |
| SimSiam | 0.952<br>$\pm 0.018$ | 0.96<br>$\pm 0.018$ | 0.713<br>$\pm 0.009$ | 0.847<br>$\pm 0.01$ | 0.868<br>$\pm 0.016$ | 0.87<br>$\pm 0.012$ | 0.681<br>$\pm 0.023$ | 0.797<br>$\pm 0.017$ |
| NNCLR | 0.984<br>$\pm 0.003$ | 0.991<br>$\pm 0.001$ | 0.737<br>$\pm 0.017$ | 0.85<br>$\pm 0.005$ | 0.916<br>$\pm 0.021$ | 0.91<br>$\pm 0.024$ | 0.709<br>$\pm 0.021$ | 0.804<br>$\pm 0.012$ |
| BYOL | 0.969<br>$\pm 0.004$ | 0.978<br>$\pm 0.004$ | 0.798<br>$\pm 0.014$ | 0.873<br>$\pm 0.02$ | 0.892<br>$\pm 0.008$ | 0.889<br>$\pm 0.007$ | 0.792<br>$\pm 0.013$ | 0.821<br>$\pm 0.018$ |
| VICReg | 0.982<br>$\pm 0.001$ | 0.985<br>$\pm 0.001$ | **0.874**<br>$\pm 0.004$ | **0.922**<br>$\pm 0.008$ | **0.959**<br>$\pm 0.005$ | **0.957**<br>$\pm 0.006$ | **0.883**<br>$\pm 0.006$ | **0.917**<br>$\pm 0.011$ |
| Barlow<br>Twins | 0.986<br>$\pm 0.001$ | 0.99<br>$\pm 0.001$ | 0.83<br>$\pm 0.011$ | 0.901<br>$\pm 0.007$ | 0.926<br>$\pm 0.01$ | 0.919<br>$\pm 0.011$ | 0.834<br>$\pm 0.007$ | 0.866<br>$\pm 0.006$ |
| Concerto | 0.973<br>$\pm 0.000$ | 0.982<br>$\pm 0.001$ | 0.769<br>$\pm 0.000$ | 0.904<br>$\pm 0.000$ | —<br>— | —<br>— | —<br>— | —<br>— |

**Table B4:** Missing modality prediction for methods trained with the CLEAR pipeline on multimodal datasets. We show the average Pearson correlation between the original and inferred missing modality: protein for PBMC CITE-seq, and ADT (protein abundance) for BMMC.

| Method | PBMC CITE-seq<br>Pearson Mean | BMMC<br>Pearson Mean |
| --- | --- | --- |
| SimCLR | **0.866** $\pm 0.001$ | 0.757 $\pm 0.002$ |
| MoCo | 0.856 $\pm 0.001$ | 0.721 $\pm 0.004$ |
| SimSiam | 0.859 $\pm 0.002$ | 0.748 $\pm 0.002$ |
| NNCLR | 0.861 $\pm 0.002$ | 0.751 $\pm 0.001$ |
| BYOL | 0.860 $\pm 0.000$ | 0.738 $\pm 0.002$ |
| VICReg | 0.865 $\pm 0.001$ | **0.759** $\pm 0.001$ |
| Barlow Twins | 0.864 $\pm 0.001$ | 0.755 $\pm 0.001$ |
| Concerto | 0.742 $\pm 0.006$ | 0.542 $\pm 0.001$ |

**Table B5:**
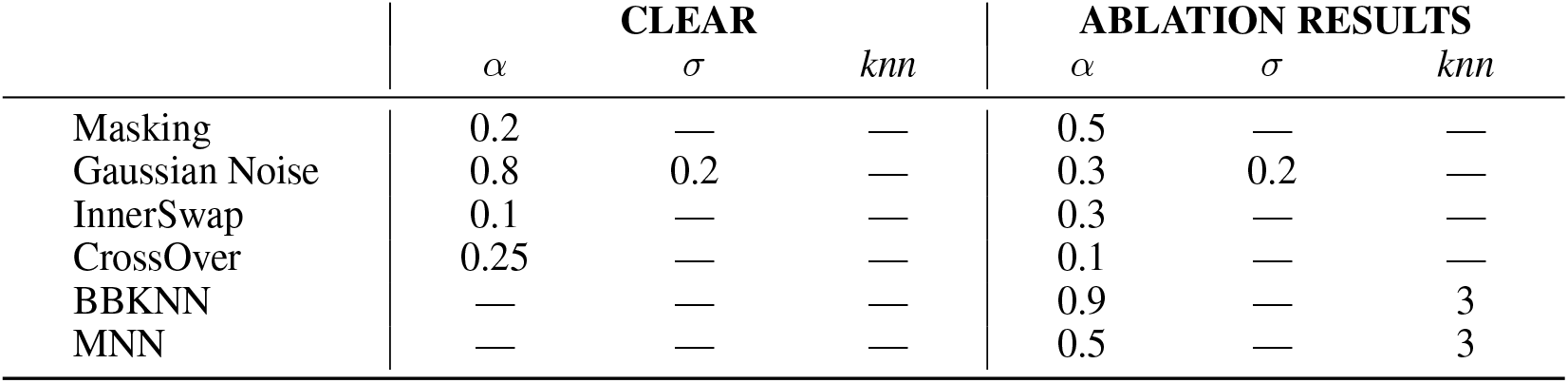
Augmentation Parameters for the CLEAR [10] augmentations on the left. Results for the ablation of all augmentations on the right, including the CLAIRE [22] augmentation denoted as MNN, and our BBKNN augmentation. Results stem from our ablation detailed in subsection A.2.

**Table B6:** Comparison of different multi-omics integration methods using the CLEAR pipeline. Data integration metrics were computed for the BMMC dataset.

| Method | Add |  |  | Concat |  |  | CLIP + Concat |  |  |
| --- | --- | --- | --- | --- | --- | --- | --- | --- | --- |
|  | Bio | Batch | Total | Bio | Batch | Total | Bio | Batch | Total |
| SimCLR | 0.827<br>$\pm 0.078$ | 0.3<br>$\pm 0.057$ | 0.616<br>$\pm 0.05$ | 0.84<br>$\pm 0.093$ | 0.273<br>$\pm 0.058$ | 0.613<br>$\pm 0.065$ | 0.511<br>$\pm 0.223$ | **0.504**<br>$\pm 0.094$ | 0.508<br>$\pm 0.166$ |
| MoCo | **0.935**<br>$\pm 0.07$ | 0.407<br>$\pm 0.019$ | **0.724**<br>$\pm 0.045$ | 0.056<br>$\pm 0.065$ | **0.8**<br>$\pm 0.000$ | 0.354<br>$\pm 0.039$ | 0.566<br>$\pm 0.157$ | 0.464<br>$\pm 0.161$ | 0.525<br>$\pm 0.049$ |
| SimSiam | 0.453<br>$\pm 0.175$ | 0.174<br>$\pm 0.025$ | 0.341<br>$\pm 0.107$ | 0.506<br>$\pm 0.146$ | 0.21<br>$\pm 0.041$ | 0.387<br>$\pm 0.083$ | 0.197<br>$\pm 0.177$ | 0.364<br>$\pm 0.051$ | 0.264<br>$\pm 0.011$ |
| NNCLR | 0.679<br>$\pm 0.171$ | 0.225<br>$\pm 0.041$ | 0.498<br>$\pm 0.113$ | 0.768<br>$\pm 0.105$ | 0.231<br>$\pm 0.034$ | 0.553<br>$\pm 0.06$ | 0.584<br>$\pm 0.088$ | 0.5<br>$\pm 0.079$ | 0.551<br>$\pm 0.066$ |
| BYOL | 0.117<br>$\pm 0.109$ | **0.8**<br>$\pm 0.000$ | 0.39<br>$\pm 0.066$ | 0.527<br>$\pm 0.029$ | 0.673<br>$\pm 0.074$ | 0.586<br>$\pm 0.03$ | 0.46<br>$\pm 0.123$ | 0.403<br>$\pm 0.152$ | 0.437<br>$\pm 0.115$ |
| VICReg | 0.791<br>$\pm 0.089$ | 0.449<br>$\pm 0.014$ | 0.654<br>$\pm 0.052$ | **0.887**<br>$\pm 0.035$ | 0.484<br>$\pm 0.009$ | **0.726**<br>$\pm 0.022$ | **0.72**<br>$\pm 0.079$ | 0.38<br>$\pm 0.055$ | **0.584**<br>$\pm 0.056$ |
| Barlow<br>Twins | 0.717<br>$\pm 0.055$ | 0.262<br>$\pm 0.026$ | 0.535<br>$\pm 0.039$ | 0.852<br>$\pm 0.096$ | 0.27<br>$\pm 0.012$ | 0.62<br>$\pm 0.059$ | 0.706<br>$\pm 0.115$ | 0.352<br>$\pm 0.085$ | 0.565<br>$\pm 0.055$ |

**Table B7:**
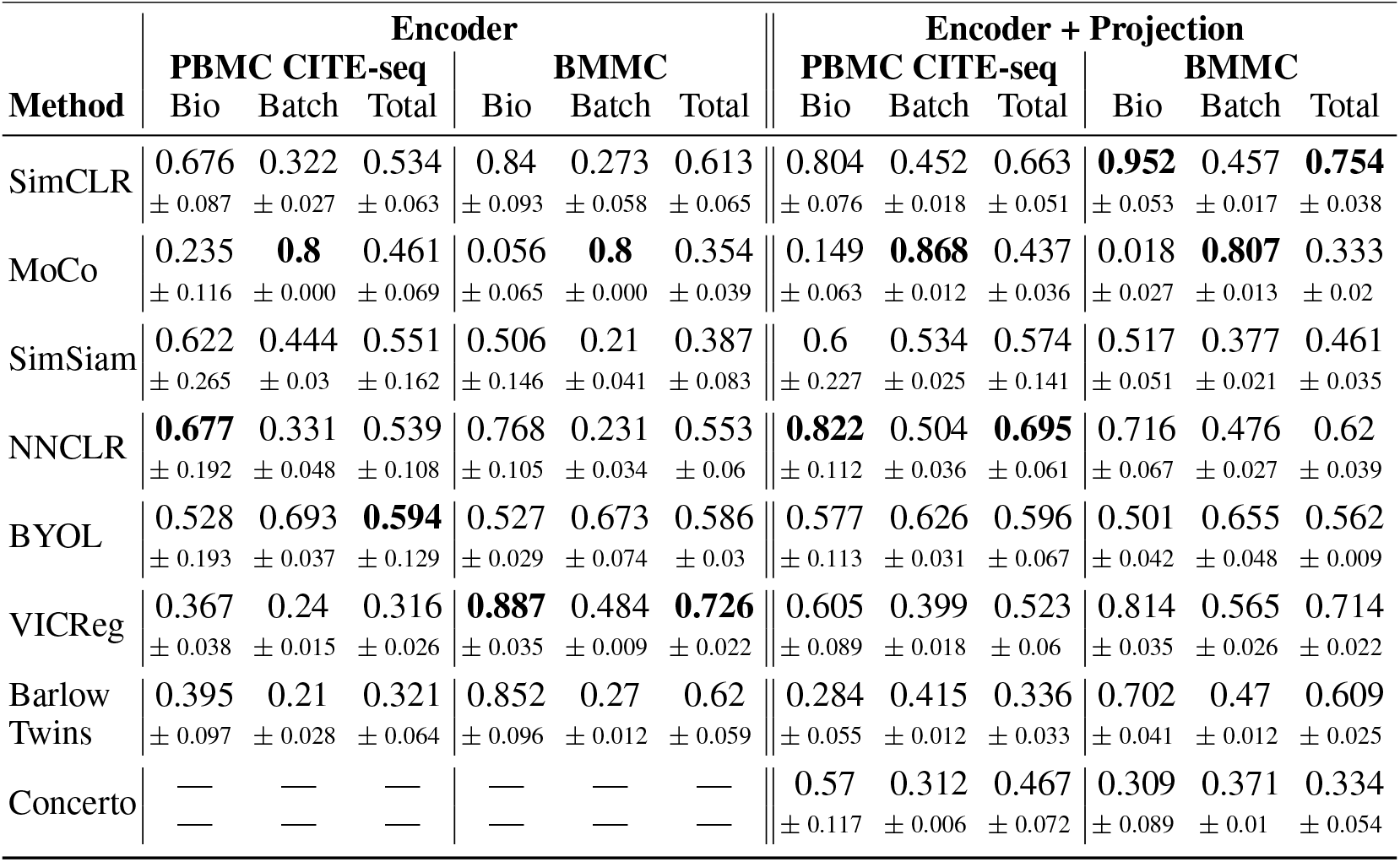
Data integration for methods trained with the CLEAR pipeline on multimodal datasets. We compare the effect of retaining the projection head during inference to the representation quality when using only the encoder.

| Method | Encoder |  |  |  |  |  | Encoder + Projection |  |  |  |  |  |
| --- | --- | --- | --- | --- | --- | --- | --- | --- | --- | --- | --- | --- |
|  | PBMC CITE-seq |  |  | BMMC |  |  | PBMC CITE-seq |  |  | BMMC |  |  |
|  | Bio | Batch | Total | Bio | Batch | Total | Bio | Batch | Total | Bio | Batch | Total |
| SimCLR | 0.676<br>$\pm 0.087$ | 0.322<br>$\pm 0.027$ | 0.534<br>$\pm 0.063$ | 0.84<br>$\pm 0.093$ | 0.273<br>$\pm 0.058$ | 0.613<br>$\pm 0.065$ | 0.804<br>$\pm 0.076$ | 0.452<br>$\pm 0.018$ | 0.663<br>$\pm 0.051$ | **0.952**<br>$\pm 0.053$ | 0.457<br>$\pm 0.017$ | **0.754**<br>$\pm 0.038$ |
| MoCo | 0.235<br>$\pm 0.116$ | **0.8**<br>$\pm 0.000$ | 0.461<br>$\pm 0.069$ | 0.056<br>$\pm 0.065$ | **0.8**<br>$\pm 0.000$ | 0.354<br>$\pm 0.039$ | 0.149<br>$\pm 0.063$ | **0.868**<br>$\pm 0.012$ | 0.437<br>$\pm 0.036$ | 0.018<br>$\pm 0.027$ | **0.807**<br>$\pm 0.013$ | 0.333<br>$\pm 0.02$ |
| SimSiam | 0.622<br>$\pm 0.265$ | 0.444<br>$\pm 0.03$ | 0.551<br>$\pm 0.162$ | 0.506<br>$\pm 0.146$ | 0.21<br>$\pm 0.041$ | 0.387<br>$\pm 0.083$ | 0.6<br>$\pm 0.227$ | 0.534<br>$\pm 0.025$ | 0.574<br>$\pm 0.141$ | 0.517<br>$\pm 0.051$ | 0.377<br>$\pm 0.021$ | 0.461<br>$\pm 0.035$ |
| NNCLR | **0.677**<br>$\pm 0.192$ | 0.331<br>$\pm 0.048$ | 0.539<br>$\pm 0.108$ | 0.768<br>$\pm 0.105$ | 0.231<br>$\pm 0.034$ | 0.553<br>$\pm 0.06$ | **0.822**<br>$\pm 0.112$ | 0.504<br>$\pm 0.036$ | **0.695**<br>$\pm 0.061$ | 0.716<br>$\pm 0.067$ | 0.476<br>$\pm 0.027$ | 0.62<br>$\pm 0.039$ |
| BYOL | 0.528<br>$\pm 0.193$ | 0.693<br>$\pm 0.037$ | **0.594**<br>$\pm 0.129$ | 0.527<br>$\pm 0.029$ | 0.673<br>$\pm 0.074$ | 0.586<br>$\pm 0.03$ | 0.577<br>$\pm 0.113$ | 0.626<br>$\pm 0.031$ | 0.596<br>$\pm 0.067$ | 0.501<br>$\pm 0.042$ | 0.655<br>$\pm 0.048$ | 0.562<br>$\pm 0.009$ |
| VICReg | 0.367<br>$\pm 0.038$ | 0.24<br>$\pm 0.015$ | 0.316<br>$\pm 0.026$ | **0.887**<br>$\pm 0.035$ | 0.484<br>$\pm 0.009$ | **0.726**<br>$\pm 0.022$ | 0.605<br>$\pm 0.089$ | 0.399<br>$\pm 0.018$ | 0.523<br>$\pm 0.06$ | 0.814<br>$\pm 0.035$ | 0.565<br>$\pm 0.026$ | 0.714<br>$\pm 0.022$ |
| Barlow<br>Twins | 0.395<br>$\pm 0.097$ | 0.21<br>$\pm 0.028$ | 0.321<br>$\pm 0.064$ | 0.852<br>$\pm 0.096$ | 0.27<br>$\pm 0.012$ | 0.62<br>$\pm 0.059$ | 0.284<br>$\pm 0.055$ | 0.415<br>$\pm 0.012$ | 0.336<br>$\pm 0.033$ | 0.702<br>$\pm 0.041$ | 0.47<br>$\pm 0.012$ | 0.609<br>$\pm 0.025$ |
| Concerto | —<br>— | —<br>— | —<br>— | —<br>— | —<br>— | —<br>— | 0.57<br>$\pm 0.117$ | 0.312<br>$\pm 0.006$ | 0.467<br>$\pm 0.072$ | 0.309<br>$\pm 0.089$ | 0.371<br>$\pm 0.01$ | 0.334<br>$\pm 0.054$ |

**Table B8:** Batch correction benchmark for methods trained using the CLEAR pipeline. This table is an extension to Table 1, containing datasets that were used during hyperparameter tuning.

| Method | HIC |  |  | MCA |  |  |
| --- | --- | --- | --- | --- | --- | --- |
|  | Bio | Batch | Total | Bio | Batch | Total |
| SimCLR | **0.879**<br>$\pm 0.02$ | 0.533<br>$\pm 0.017$ | 0.741<br>$\pm 0.014$ | **0.919**<br>$\pm 0.02$ | 0.557<br>$\pm 0.04$ | **0.774**<br>$\pm 0.011$ |
| MoCo | 0.831<br>$\pm 0.012$ | 0.638<br>$\pm 0.017$ | 0.754<br>$\pm 0.01$ | 0.316<br>$\pm 0.071$ | **0.86**<br>$\pm 0.033$ | 0.534<br>$\pm 0.04$ |
| SimSiam | 0.606<br>$\pm 0.039$ | 0.466<br>$\pm 0.068$ | 0.55<br>$\pm 0.039$ | 0.358<br>$\pm 0.073$ | 0.592<br>$\pm 0.022$ | 0.452<br>$\pm 0.044$ |
| NNCLR | 0.738<br>$\pm 0.022$ | 0.45<br>$\pm 0.012$ | 0.623<br>$\pm 0.011$ | 0.604<br>$\pm 0.1$ | 0.506<br>$\pm 0.024$ | 0.565<br>$\pm 0.059$ |
| BYOL | 0.56<br>$\pm 0.021$ | **0.863**<br>$\pm 0.034$ | 0.681<br>$\pm 0.024$ | 0.156<br>$\pm 0.089$ | 0.695<br>$\pm 0.116$ | 0.372<br>$\pm 0.095$ |
| VICReg | 0.881<br>$\pm 0.038$ | 0.647<br>$\pm 0.012$ | **0.787**<br>$\pm 0.024$ | 0.829<br>$\pm 0.046$ | 0.642<br>$\pm 0.032$ | 0.755<br>$\pm 0.029$ |
| Barlow<br>Twins | 0.863<br>$\pm 0.009$ | 0.534<br>$\pm 0.007$ | 0.732<br>$\pm 0.004$ | 0.804<br>$\pm 0.048$ | 0.53<br>$\pm 0.025$ | 0.694<br>$\pm 0.026$ |
| Concerto | 0.0<br>$\pm 0.0$ | 0.508<br>$\pm 0.003$ | 0.204<br>$\pm 0.001$ | 0.808<br>$\pm 0.009$ | 0.383<br>$\pm 0.0$ | 0.638<br>$\pm 0.005$ |

